# Step-wise Hydration of Magnesium by Four Water Molecules Precedes Phosphate Release in a Myosin Motor

**DOI:** 10.1101/817254

**Authors:** M.L. Mugnai, D. Thirumalai

## Abstract

Molecular motors, such as myosin, kinesin, and dynein, convert the energy released by the hydrolysis of ATP into mechanical work, which allows them to undergo directional motion on cytoskeletal tracks. This process is achieved through synchronization between the catalytic activity of the motor and the associated changes in its conformation. A pivotal step in the chemomechanical transduction in myosin motors occurs after they bind to the actin filament, which triggers the release of phosphate (P_i_, product of ATP hydrolysis) and the rotation of the lever arm. Here, we investigate the mechanism of phosphate release in myosin VI, which has been debated for over two decades, using extensive molecular dynamics simulations involving multiple trajectories each several *μs* long. Because the escape of phosphate is expected to occur on time-scales on the order of milliseconds in myosin VI, we observed P_i_ release only if the trajectories were initiated with a rotated phosphate inside the nucleotide binding pocket. The rotation provided the needed perturbation that enabled successful expulsions of P_i_ in several trajectories. Analyses of these trajectories lead to a robust mechanism of P_i_ release in the class of motors belonging to the myosin super family. We discovered that although P_i_ populates the traditional “back door” route, phosphate exits through various other gateways, thus establishing the heterogeneity in the escape routes. Remarkably, we observe that the release of phosphate is preceded by a step-wise hydration of the ADP-bound magnesium ion. In particular, the release of the anion occurred *only after four water molecules* hydrate the cation (Mg^2+^). By performing comparative structural analyses, we suggest that the hydration of magnesium is the key step in the phosphate release in a number of ATPases and GTPases that share a similar structure in the nucleotide binding pocket. Thus, nature may have evolved hydration of Mg^2+^ by discrete water molecules as a general molecular switch for P_i_ release, which is a universal step in the catalytic cycle of many machines which share little sequence or structural similarity.

## Introduction

Myosins are molecular motors that, fueled by the hydrolysis of ATP, move unidirectionally on a cytoskeletal track (the filamentous actin, or F-actin) (1, 2). In order to perform their functions, myosins, like other motors, undergo a chemomechanical cycle in which the nature of the nucleotide bound to the enzyme [no nucleotide (apo), ATP, ADP and P_i_, and ADP] is coupled with the three-dimensional structure attained by the myosin head, and with nucleotide-dependent affinity for actin (Fig. 1a). After catalyzing the hydrolysis of ATP, myosin binds to the filament; this triggers the release of phosphate and the “power-stroke,” a conformational transition in which the movement of the converter domain (black in Fig. 1a) is amplified by the rotation of the lever arm (cyan in Fig. 1a, where only a small fragment of the full lever arm is shown). While myosin ADP-bound and apo states are tightly bound to actin, the binding of ATP induces the dissociation of the actomyosin complex. Finally, the hydrolysis of ATP is coupled to the “re-priming” stroke, a transition during which the lever resumes the pre-power-stroke conformation, thereby preparing myosin for a new cycle.

**Figure 1:**
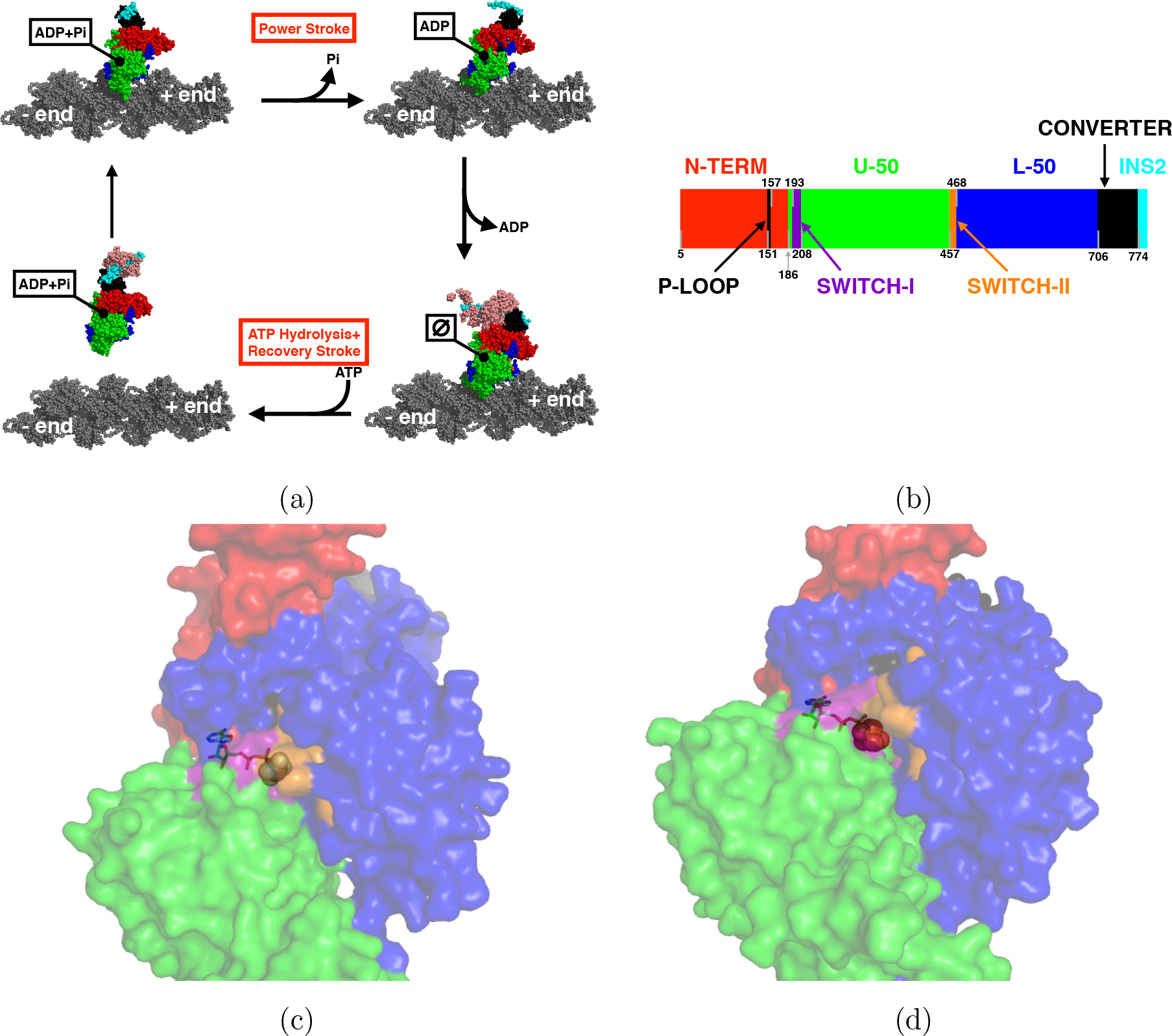
Sequence, cycle, and structure of myosin VI. (a) Myosin cycle. The different conformations are taken from a combination of structural studies – actin and ADP-bound myosin (top right) [PDB 6BNW (44)], rigor (or apo, bottom right) [2BKI (45)], pre-power stroke (ADP+P_i_, detached from actin, bottom left) [4ANJ (26), crystallized with a phosphate analog], and phosphate release (P_i_R, ADP-P_i_, attached to actin, top left) [4PJM (6)]. Only the structure of the ADP-bound state was solved in the actin-bound conformation; in the other cases, the myosin was aligned to the ADP-bound myosin. (b) Sequence of myosin VI. The color code for representing the different structural elements is kept throughout the paper. The identification of individual elements follows indications from multiple sources (1, 2, 6). However, we do not show some groups that we do not discuss in the paper (for instance, the relay helix) and the domains that we consider do not correspond precisely to the proteolytic fragments (1). (c) Filled-mode representation of myosin VI in PrePS [PDB 4ANJ (26)]. ADP and P_i_ are shown as sticks, and can be seen through the transparent motor. The “back door” pathway is closed in the PrePS conformation (see the purple orange shades showing Switch I and II). (d) The P_i_R conformation is represented in filled mode, with the ligands shown as sticks [PDB 4PJM (6)]. The “back door” pathway is now open – the purple and orange shades (Switch I and II) have moved away from each other and the phosphate is more clearly visible.

The pivotal step of the chemomechanical transduction is the power stroke, which occurs when myosin binds to the actin filament after ATP hydrolysis. The interaction between the motor and the track is believed to stabilize a conformation which accelerates phosphate release [see (3) for myosin VI] and produces the forward swing of the lever arm. In order to elucidate the details of this crucial step in the myosin catalytic cycle, it is therefore necessary to achieve a detailed molecular-level understanding of the processes governing phosphate release, which is possible using atomically detailed molecular dynamics simulations.

When the first structure of a myosin bound to ADP and an analog of P_i_ was resolved (4), the release pathway of P_i_ immediately posed a puzzle. ATP binds the motor domain with the γ-phosphate buried inside the nucleotide binding site. As a consequence, it is possible that ADP hinders the release of P_i_ from the entrance gateway, suggesting that an alternative exit route must exist. Yount *et al.* postulated that there is a “back door” pathway for the release of P_i_ along the seam that separates the so-called U-50 and L-50 domains of the motor (5) (see Fig. 1). Although such a release route appears to be open when the nucleotides are missing from the binding site, in the pre-power-stroke (PrePS) conformation – crystallized with ADP and an analog of P_i_ bound to myosin – this route is closed by two loops, Switch-I and Switch-II (Fig. 1c). Thus, either or both the loops must move to open the “back door” channel, although such displacements are also expected to reduce the efficiency of the motor (6). A solution to this puzzle emerged when the crystal structure of a myosin in a putative P_i_-release (P_i_R) conformation was resolved (6). The structure shows partial movement of Switch-II which opens the “back door” pathway (Fig. 1d), and a change in the actin-binding interface that was suggested to strengthen the interaction with F-actin (6). In addition, the converter is found in prestroke position, indicating that P_i_ release precedes the power stroke, and a combination of kinetics and structural experiments supported the interpretation of this structure as the P_i_R conformation (6). These findings are in agreement with optical trap experiments for myosin V (7), although at odds with experiments in muscle fibers that identified the existence of a force-generating state prior to phosphate release (8).

Although it appears that the “back door” pathway for P_i_ release is open in the P_i_R conformation, this might not be the unique exit route for P_i_, as suggested by computational studies adopting enhanced sampling techniques (9). Furthermore, a clear picture of the microscopic conformational transitions leading to the P_i_ release is lacking. In particular, the role that water molecules play in enabling P_i_ release has not been probed. In order to remedy this situation, we investigate the mechanism of P_i_ release from myosin VI (MVI) nucleotide binding site by running many multi-microsecond Molecular Dynamics (MD) trajectories generated using the supercomputer Anton 2 (10). Experimentally, the actin-activated P_i_ release time from the PrePS conformation is expected to be in the range 8.3ms-33ms (3, 11, 12), which is still beyond the current computational capabilities for conventional all-atom MD simulations of systems as large as myosin [although enhanced sampling methods, for instance (9, 13), and spatial (14) or temporal (15) coarse-graining have allowed to access longer timescales]. Accordingly, we only observed the release of phosphate within the timescale accessible to Anton 2 when the simulations were started with a rotated P_i_, which we believe is sufficient to reveal the complexities of P_i_ release. This perturbation weakens the interaction between the phosphate and the ADP-bound Mg^2+^ because a P_i_ hydroxyl group (instead of an oxygen of P_i_) is placed in the coordination shell of the cation. In what follows we refer to “CA” (Crystallographically Aligned) as the initial conformation attained by ensuring that a phosphate oxygen group points towards the magnesium ion (see Fig. 2b); we term instead “RA” (Rotated Alignment) the structure in which the phosphate is initially rotated (see Fig. 2d).

**Figure 2:**
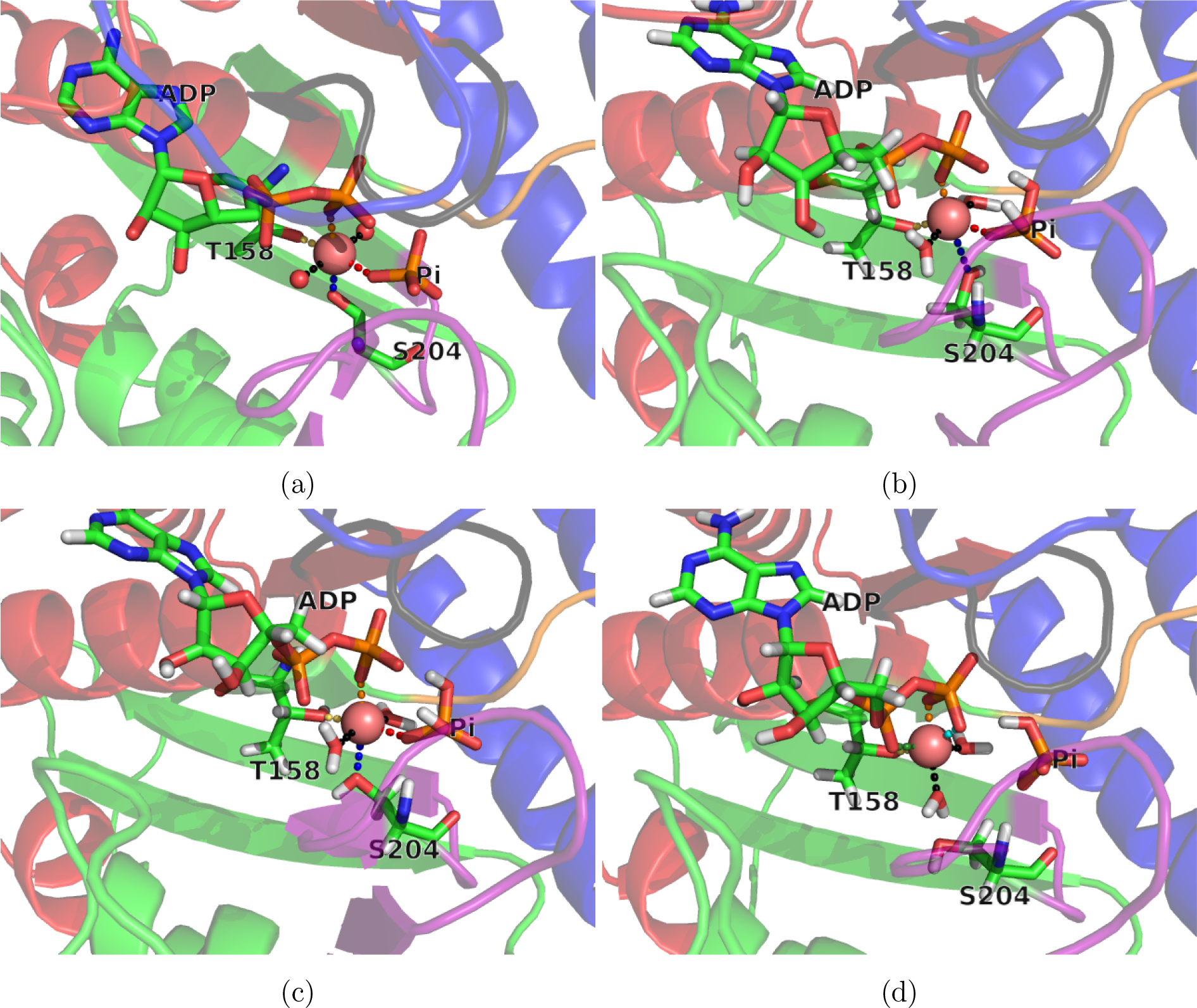
Coordination of Mg^2+^ after equilibration. (a) In the crystal structure (4PJM) Mg^2+^ ion is coordinated by two water molecules (shown as spheres), T158, S204, Pi, and one oxygen from ADP *β*-phosphate. (b) Conformation of the nucleotide binding site after equilibration starting with the CA initial conformation. Similarly to the crystal structure (a), the first shell of Mg^2+^ is occupied by T158, S204, P_i_, and one oxygen from ADP *β*-phosphate. (c) Initial structure for the simulations carried out with the 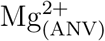 divalent cation. The coordination shell of Mg^2+^ is not affected by the choice of the force field. (d) Nucleotide binding pocket of MVI after equilibration with the P_i_ rotated to point a hydroxyl group towards Mg^2+^ (RA initial conformation). Distortion induced by the different orientation of the P_i_ modifies the first shell of Mg^2+^, which is now made by two water molecules, three oxygens from ADP diphosphate, and T158.

Strikingly, we found that water molecules play a crucial role in the escape of phosphate. First, we observed that the magnesium cation is progressively hydrated in a step-wise manner. Second, in all the simulations resulting in the release of phosphate, before the P_i_ leaves the proximity of ADP four water molecules hydrate the magnesium, upwards from the two found in the crystal structure. This leads us to suggest that the hydration of magnesium by four discrete water molecules, which serves as a switch, is the step that triggers the release of phosphate. By comparing the structures of the nucleotide binding pockets in other biological machines, we believe that this important discovery might constitute a general mechanism for phosphate release in many families of ATPases and GTPases [for instance myosins, kinesins, and G-proteins (16)]. We show that these machines share similar structural features in their nucleotide binding sites. Thus, we have discovered using computations a unifying mechanism of P_i_ release in a class of biological machines. Our findings are amenable to experimental tests.

## Results

### Analysis of the initial structure

In Fig. 2a we show the first coordination shell of magnesium in the crystal structure. Two water molecules, an oxygen from ADP *β*-phosphate, and P_i_ coordinate Mg^2+^ together with the side chains of T158 and S204 (we use one letter code for amino acids). The same first coordination shell of Mg^2+^ is found after the initial relaxation from the CA conformation (Fig. 2b). The use of an optimized force field for magnesium (17) (referred to as 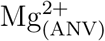) did not change this observation (see Fig. 2c). In agreement with quantum mechanical simulations (18), one hydroxyl group of P_i_ is directed towards the *β* phosphate of ADP, the other is in contact with the side chain of S203. According to a more recent study in myosin (19), the hydroxyl group of P_i_ close to the *β* phosphate is stabilized by a hydrogen bond with the side chain of a serine in the P-loop (S153 in MVI). In the 4PFP structure (and choosing conformation A of 4PJM) the side chain of S153 is close to the hydroxyl group of P_i_, which in turn is in proximity to S203. In our case, the hydroxyl group of the side chain of S153 points away from the phosphate due to the choice of one of the initial rotamers (B) present in the crystal structure. The presence of this hydrogen bond might provide additional stabilization of P_i_ in the binding pocket. Thus, it is possible that its absence might help the release of P_i_.

After a short relaxation of the initial RA structure, the nucleotide binding pocket is distorted: an oxygen from ADP *α*-phosphate and one from ADP *β*-phosphate have replaced P_i_ and S204 in the first coordination shell of Mg^2+^ (see Fig. 2d). It is reasonable to suggest that this conformational change weakens the interaction between the anion (P_i_) and the cation (Mg^2+^).

### Coordination of Mg^2+^ and P_i_ release: simulations starting from CA conformation

In all the simulations beginning from the CA conformation, regardless of the force field used for Mg^2+^, we do not observe P_i_ release on the time scales of the simulations, which are several microseconds. Comparing Fig. 2b and Fig. 3b, it is clear that the side chains of T158 and S204 are replaced in the Mg^2+^ coordination shell by an oxygen of the ADP and an extra water molecule. As shown in Fig. 3a (and Fig.s S2-S3), this exchange occurs within the first microsecond and it remains stable for the rest of the simulation. In most trajectories, three water molecules hydrate Mg^2+^. In one simulation, carried out using 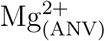, four water molecules surround the cation for short time intervals, before the hydration observed in the crystal structure is restored, thus preventing P_i_ release. Notably, lowering the charge of the magnesium to 1.5 did not result in the expulsion of P_i_ within a few microseconds (see Fig. S5).

**Figure 3:**
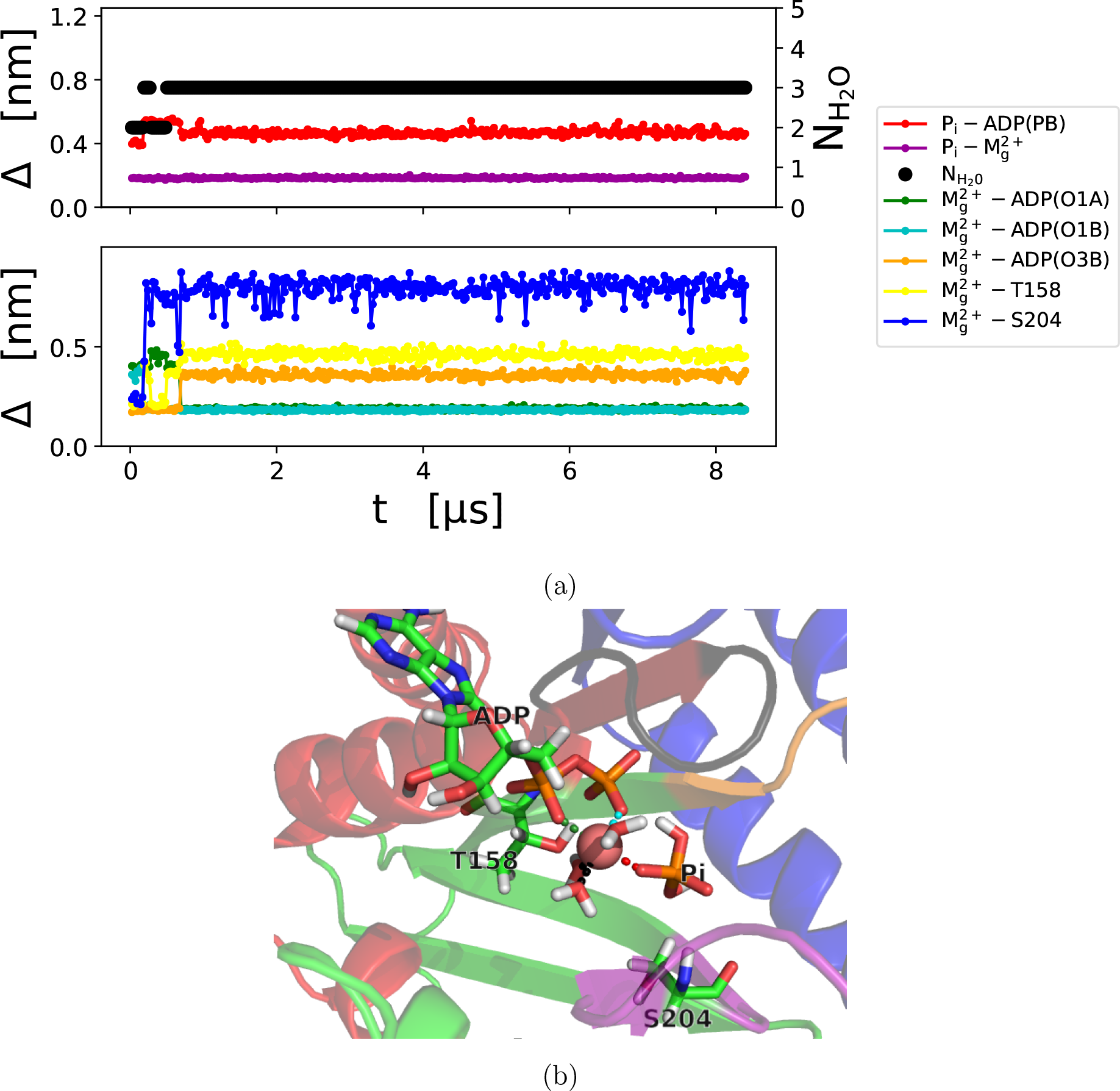
Conformation of the nucleotide binding site during a simulation carried out from the CA initial conformation with the CHARMM energy function. (a) Time-dependent distances of atoms engaged in the first solvation shell of Mg^2+^. The color code is given in the legend. Red indicates the distance between the *β* phosphorous of ADP and P_i_. The distances between the oxygen of ADP *α* (*β*) phosphate and Mg^2+^is in green (light blue and orange). The contacts between the side chains of the residues T158 and S204 with Mg^2+^are shown in yellow and blue, respectively. The purple line shows the distance between Mg^2+^ and the oxygen of P_i_ which is closest to the cation. Finally, the black dots indicate the number of water molecules in the first shell of Mg^2+^, computed by counting the number of water oxygens within 2.75Å from Mg^2+^. The x-axis shows time in *μ*s, the left y-axis the distance in nm, and the right y-axis the number of water molecules in the first solvation shell of Mg^2+^. (b) The conformation of the nucleotide binding site after approximately 2.4*μ*s.

### Coordination of Mg^2+^ and P_i_ release: simulations starting from the RA conformation

The P_i_ is released in 6 out of the 10 simulations generated starting from the RA initial condition. Remarkably, in all of these trajectories, the escape of P_i_ from the nucleotide binding site occurs only after a fourth water molecule occupies the first solvation shell of Mg^2+^ (see Fig. 4 and Fig. S4), normally displacing an oxygen of ADP diphosphate group, and in one case the side chain of T158. Note that in one trajectory the P_i_ re-enters the nucleotide binding site towards the end of the run. In all the other simulations, once the phosphate is released it remains solvated (see Fig. S4). These observations allow us to dissect the pathways of P_i_ release, which we discuss below.

**Figure 4:**
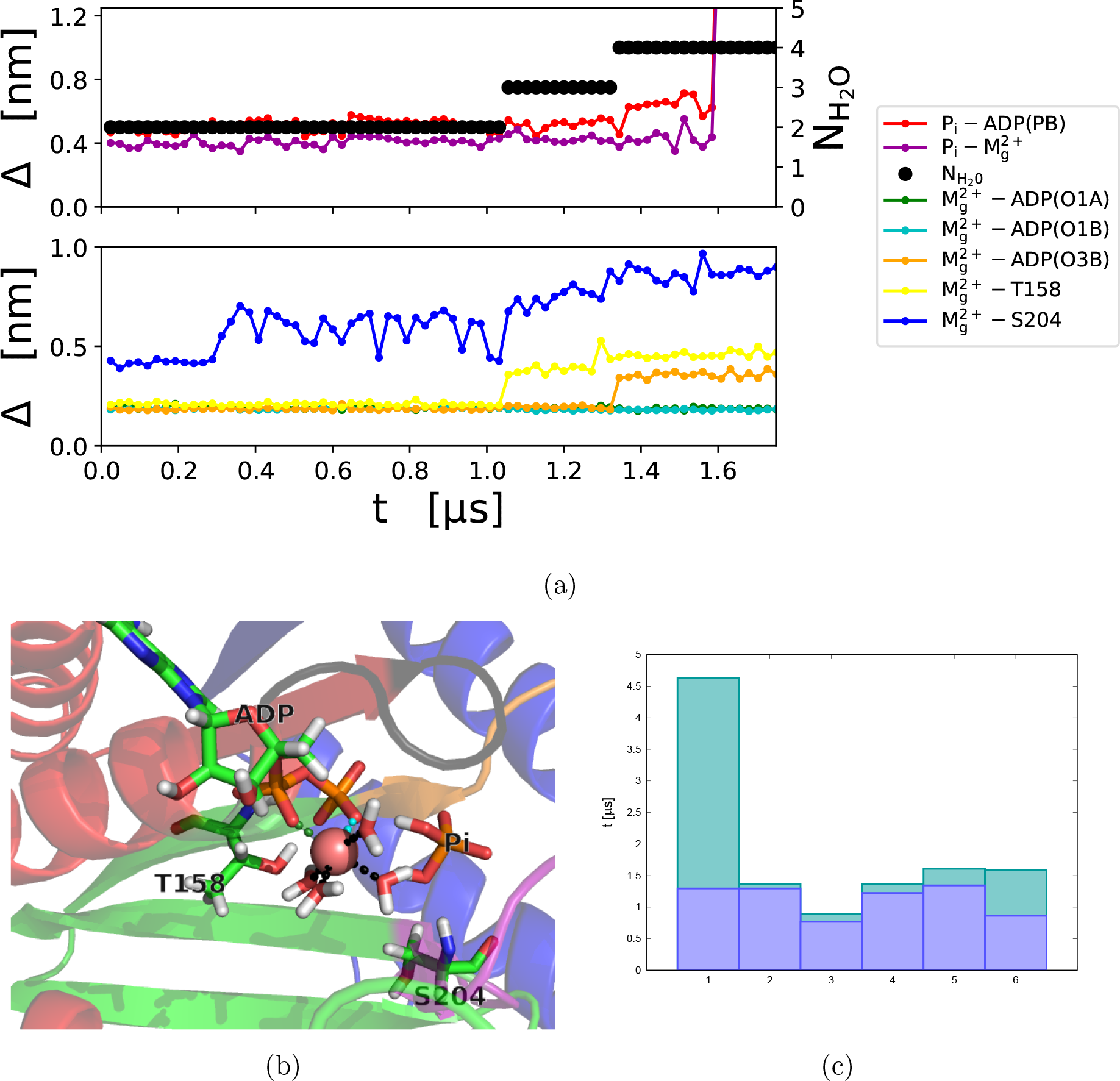
Conformation of the nucleotide binding site during simulations carried out from the RA initial conformation. (a) Time-dependent distances of atoms engaged in the first solvation shell of Mg^2+^. The color code (reported in the legend) is the same as in Fig. 3a. The trajectory is shown up to the the release of phosphate; the entire trajectory is reported in the Supplementary Information (see Fig. S4g), and shows the re-entry of the phosphate. (b) The conformation of the nucleotide binding site after approximately 1.3*μ*s, when the fourth water molecule joins the first coordination shell of Mg^2+^. (c) Time for the stable arrival of the fourth water (blue) and for release of the phosphate (green). The conformations are sampled with a time resolution of 24ns.

In one simulation (see Fig. S4h) the binding of a fourth water molecule to Mg^2+^ does not induce the release of phosphate and it is followed by the rotation of P_i_ to the CA conformation with three water molecules in the first coordination shell of the cation. This indicates that (i) the arrival of the fourth water molecule is not sufficient to induce the expulsion of the phosphate, even in the RA conformation; (ii) the arrival of the fourth water molecule is reversible; (iii) it is possible to undergo a RA→CA transition. Finally, we also report a RA→CA transition during the equilibration phase of a simulation set up in the RA conformation with the ANV magnesium.

### Four different exit pathways

We identified four channels through which the P_i_ can be released. In two trajectories, the phosphate leaves from the expected backdoor pathway (purple circle in Fig. 5), although it can also escape through the top (red circles in Fig. 5) or side route (orange circle in Fig. 5). Finally, in one case the phosphate leaves through the front channel, nearby the ADP (highlighted by a light-blue circle in Fig. 5). We examined also the location of P_i_ during the escape (Fig. 6a-6b) and compared it with (i) the position of the phosphate during the trajectories that show the release but before the fourth water molecule hydrates Mg^2+^(Fig. 6c), and (ii) with the simulations initiated in the CA conformation (Fig. 6d-6e). Before the release, P_i_ is confined in the vicinity of ADP, though in one trajectory we found that the position is distorted and the phosphate moves close to the front exit channel (Fig. 6d). On the other hand, during the escape P_i_ predominantly populates the expected pathway (Fig. 6a-b), in agreement with the crystal structures of myosin VI in the PiR state after they were soaked in a solution containing large quantities of phosphate (6), which shows two P_i_, one by the exit of the tunnel and another in the N-terminal subdomain.

**Figure 5:**
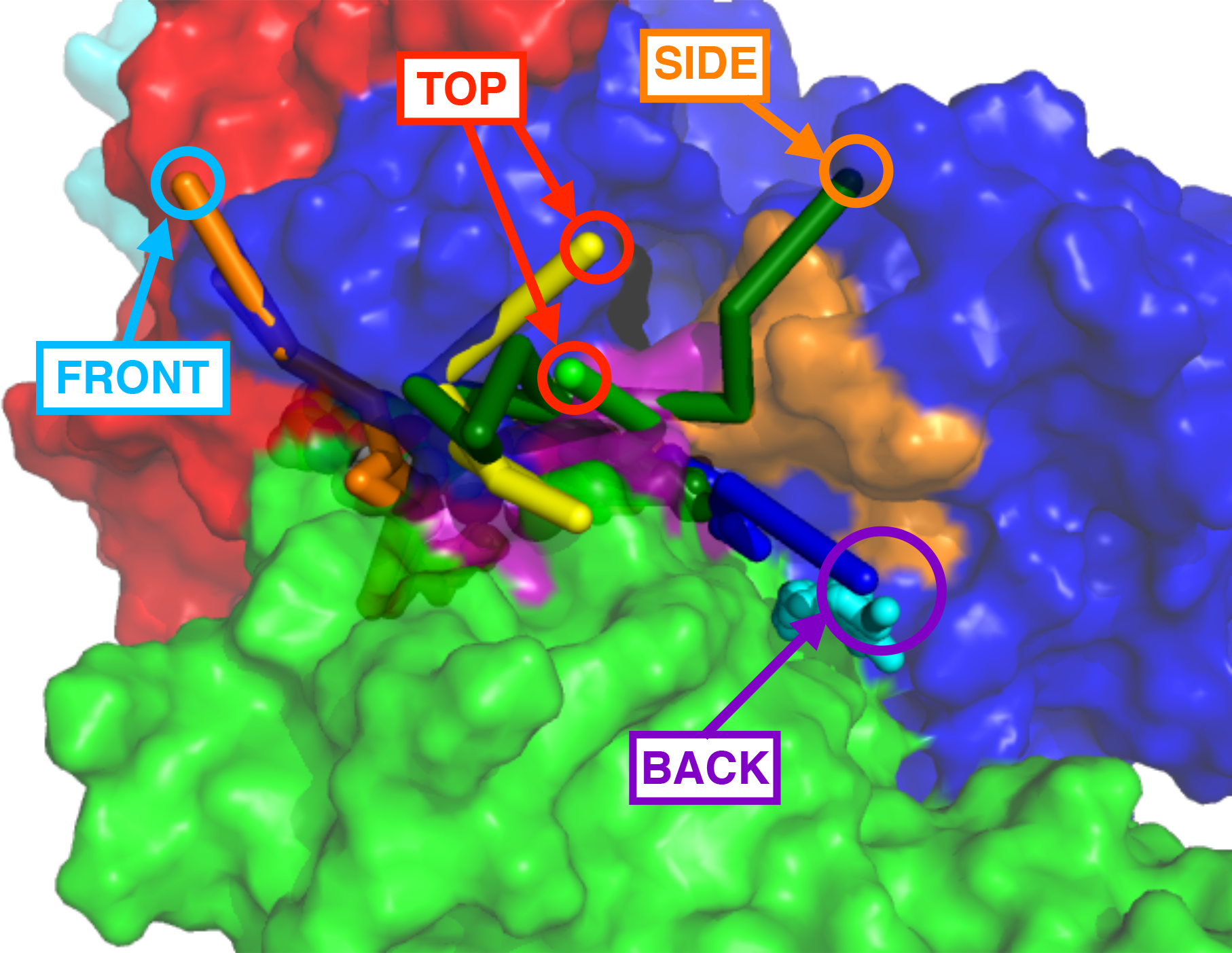
Trajectories of P_i_ release. Two trajectories are released from the back door (see purple circle). In the red and orange circles are highlighted the trajectories that are released from the top or the side of the backdoor pathway, respectively. The only trajectory released from the front path - by the ADP - is highlighted with a cyan circle.

**Figure 6:**
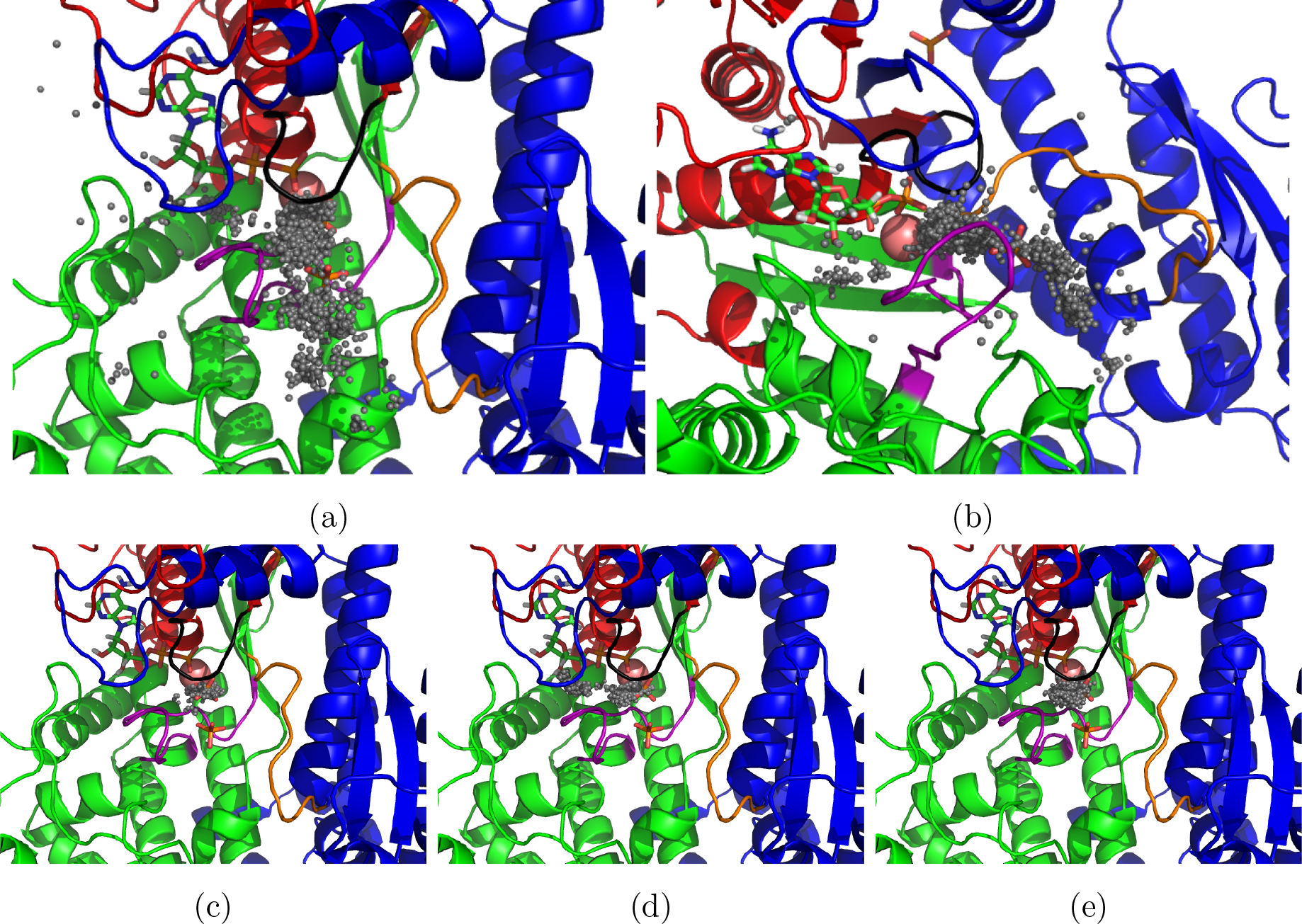
Location of P_i_. The grey dots indicate the position of the phosphorous atom of P_i_. The conformations sampled during the different trajectories were aligned to a reference structure, of which we report the protein (cartoon), ADP (sticks), and Mg^2+^ (red sphere). We report as sticks also the location of the P_i_ identified in structure 4PFP and after short soaking of the crystal in a phosphate bath (4PJN) (6), which shows P_i_ at the exit of the tunnel and in another location in the L-50 domain. (a-b) Trajectories during the last ≈120ns before the release of P_i_, showing only conformations during which four water molecules are bound to the cation. The different points shown are taken from the six trajectories showing the release of P_i_ at time resolution of 240ps. The time for the escape was identified from the first frame showing the distance between the phosphorous of P_i_ and the *β* phosphorous of ADP exceeded 1.5nm with a resolution of one frame per 24ns. Panel (a) is a top view, the perspective from a side is shown in panel (b). (c) Location of P_i_ during the trajectories starting in the RA conformation showing phosphate release, but with three molecules bound to Mg^2+^, therefore before the escape of phosphate. Points are sampled at a time resolution of 24ns. (d-e) Position of the phosphate during the simulations begun in the CA conformation and carried out using CHARMM Mg^2+^ or 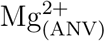 (e) - time resolution: 24ns.

### Flexibility of the converter and the N-terminal domain

The converter in myosin VI is pliant: in most trajectories, regardless of the setup, it fluctuates away from the crystal-structure pose (see Fig. S7) without significantly changing its conformation (see lower figure in each panel of Fig. S7). We also note that the converter in the simulations initiated with the rotated P_i_ seemed to move the converter earlier than those with the phosphate in the correct orientation. This might suggest some communication between the nucleotide binding site and the converter. Surprisingly, we found that in a few simulations the N-terminal *β*–barrel (NTBB) is mobile (see Fig. S8). As for the converter, the structure of the NTBB does not change significantly (see lower figure of each panel in Fig. S8), but the whole system detaches from the motor domain and in some cases finds its position close to the converter.

### Movement of the ADP

We monitor the conformation of the ADP sugar as the distance between a nitrogen of ADP (N6) and the alpha carbon of residue N98. Looking at the distribution of distances, we set a threshold at 0.725nm: if the distance exceeds this threshold the nucleoside of ADP is deemed to be outside of its binding site. As shown in Fig. S9, ADP flips out of the pocket more commonly when we start with the rotated P_i_ and while the phosphate is still bound. Strikingly, after the P_i_ has left the pocket, the ADP nucleoside maintains its contact with the protein, thus indicating that the rotated P_i_ might introduce some “stress” in the binding pocket. Interestingly, in the simulations starting with the CA conformation we occasionally see that the ADP sugar flips out of the pocket only when the CHARMM potential for Mg^2+^ is used, although the statistics is likely not sufficient to make conclusive statements.

## Discussion

### Water triggers the escape ofP_i_, and prepares ADP for the release

As illustrated in Fig. 4, Fig. 6 and Fig. S4 the phosphate does not move away from Mg^2+^ADP until four water molecule hydrate the cation. This is the major finding of our simulations. It is reasonable that the arrival of a fourth water molecule may weaken the interaction between the cation (Mg^2+^) and the anion (P_i_) by screening the electrostatic attraction and by sterically obstructing the position of the phosphate. In addition, indirect effects could also play a role. For instance, the arrival of the fourth water molecule in the coordination shell of Mg^2+^ requires re-arrangement of the ADP di-phosphate group, which may in turn affect the interaction with P_i_.

In some trajectories there is a long waiting time between the arrival of the fourth water molecule and the release of phosphate (see Fig. 4a and Fig. S4). We also observed instances in which a transitory four-water-molecule hydration of P_i_ does not lead to the escape of phosphate, or trajectories in which P_i_ is mobile although only three water molecules bind the cation. This indicates that other interactions, which may not be directly affected by the hydration of magnesium, contribute to holding the phosphate in the vicinity of ADP. Therefore, the hydration of magnesium appears to be a necessary but may not be sufficient for phosphate release. Nevertheless, our findings show that the if four water molecules are coordinated to Mg^2+^ release of P_i_ occurs with high probability.

After the escape of phosphate, the first coordination shell of magnesium remains occupied by four water molecule and two oxygens from ADP. This is likely to weaken the binding between the protein and ADP. We surmise that not only the hydration of Mg^2+^ triggers the escape of phosphate, but it also prepares the release of the nucleotide. The

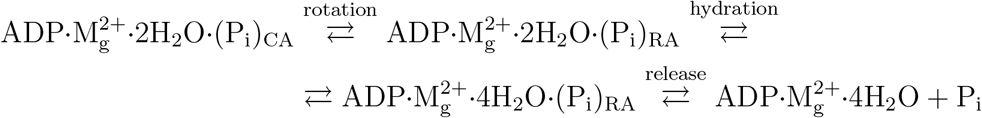

#### Scheme 1

Phosphate release mechanism in which the rotation of the phosphate is rate limiting. Note that the simulations indicate that the speculated CA→RA transition and the observed hydration of magnesium are reversible.

catalytic cycle cannot be completed until these events take place.

### Rotation of phosphate in the binding site

Studies of ATP hydrolysis-induced oxygen exchange in the γ-phosphate showed that the ATP hydrolysis reaction is reversible (20–22). Moreover, the data from quenched-flow experiments shows that the measurements are well-described by a model in which all of the oxygens of P_i_ are treated identically (22). This implies that after hydrolysis P_i_ could rotate freely inside of the binding site within the timescale of release. The presence of actin results in a reduced oxygen exchange, which is likely due to the increased rate of phosphate release in the presence of actin (22), a feature that could be justified if actin stabilizes the formation of the PiR structure.

Taking these studies into consideration, we propose the following model for phosphate release. After hydrolysis the phosphate rotates in the nucleotide binding site and transitorily orients itself in a RA conformation; then the magnesium is hydrated with four water molecules, which leads to the release of the anion (see Scheme 1). We did not observe a CA→RA transition during our simulations; however, we report two RA→CA transitions, one of which is concomitant with the reduction of the number of water molecules bound to magnesium from 4 to 3 (see Fig. S4h). This indicates that the CA→RA transition may be possible, although because we never observe it we conclude that (i) it is slower than the simulation time-scale, and (ii) RA is metastable compared to CA. Importantly, though, in the most interesting of the RA→CA transition, the rotation of the phosphate is associated with the loss of one water molecule in the coordination shell of magnesium: if the reverse transition does occur on longer timescales, it may be paired with the hydration of the cation.

Nevertheless, it is possible that the hydration of magnesium occurs in the CA conformation as well, which means that the RA conformation is not a necessary intermediate. This second case is shown in Scheme 2. In Scheme 1 attaining the RA conformation is the bottleneck for phosphate release; in Scheme 2 the hydration of the cation could be the slow process. Overall, whether the RA conformation is an intermediate facilitating P_i_ escape remains a conjecture that needs to be experimentally tested.

### The escape pathways are heterogeneous

Our simulations suggest that, although during release the P_i_ populates predominantly the “back door” pathway, the escape

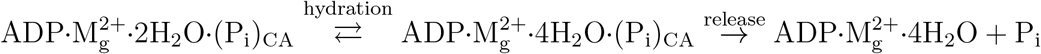

#### Scheme 2

Phosphate release mechanism in which the reversible hydration of the phosphate is rate limiting. In this scenario, the RA conformation is not an intermediate.

may occur through multiple gateways. The presence of multiple exit pathways was previously suggested using simulations based on the Locally-Enhanced Sampling (LES) method (9). Compared to straightforward simulations, LES and other enhanced sampling techniques (13) have the advantage of providing information about the probability of escaping through different channels. On the other hand, our direct simulation protocol enabled us to propose the hydration of the Mg^2+^ as a step of phosphate release; such an insight is most readily obtained via straightforward simulations.

### Assessment of Force Field for Mg^2+^

We ran simulations with two types of Mg^2+^: (i) from the CHARMM36 force field (Fig. S2), (ii) from an improved parametrization (17) (Fig. S3). When initializing the simulations in the CA conformation we did not see the release of phosphate on the time-scale of several *μs*. The initial structures of the nucleotide binding site obtained with the CHARMM36 Mg^2+^ and with 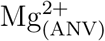 are similar, and are close to the crystal structure. In both cases though, in a few hundreds of nanoseconds the interactions between T158 and S204 and Mg^2+^ are lost and a new conformation is found in which more oxygens of ADP are in the first solvation shell of the cation. We made the same observation during a simulation performed with the alpha carbon of all residues (with the exception of T158 and S204) fixed (see Fig. S6), indicating that the breakage of the interaction between Mg^2+^ and the residues T158 and S204 is likely not due to some large conformational change in the nucleotide binding pocket, and it appears to be the arrangement in the nucleotide binding site, which is influenced by the force field. We note that the development of force fields for divalent cations is an active field [see for a recent paper (23)]; the outcome of these studies should help to increase the accuracy of the models such as the one that we discussed here. Despite potential effects of force fields, we believe that based on physical considerations, experimental findings, and comparisons with other biological machines (see below) the proposed mechanism of phosphate release is robust.

### Overall structure of the motor

Overall, the structure of the motor is intact throughout the simulations. However, as we discussed, regions such as the converter and the NTBB display a large degree of mobility. A recent experimental and numerical study of myosin VI re-priming stroke has documented a high level of flexibility for the converter, which occupies a conformation intermediate between the pre- and post-stroke orientations reported by previous structural investigations (24). Similarly, starting from the PiR, we find that the converter partially moves towards the post-stroke conformation. More surprising is the mobility of the NTBB. We recorded an unhinged NTBB for simulations starting with both CA and RA initial conformations, thus suggesting that the state of the nucleotide binding site might not be the primary cause of the movement of NTBB. We note that the crystal structure that we used was solved in the presence of glycerol and isopropyl alcohol. In addition, we did not account for the presence of the actin filament. If the PiR structure is indeed stabilized by the interaction with actin, it is possible that some unexpected modes of fluctuations occur in the structure. Further tests would be necessary in order to understand the cause and meaning of the large fluctuations of the converter and NTBB domain.

### Generality of the result

The generality of the central result that a four-fold hydration of Mg^2+^ is necessary for the release of phosphate could be assessed by examining the structures of the nucleotide binding pocket in other systems. Walker *et al.* (25) observed that many ATPases and GTPases shared common motifs, one of which - denoted as motif A - became known as the the P-loop. Once the structures for a series of ATPases and GTPases had been solved, Vale (16) pointed out that G-proteins, myosins, and kinesins are characterized by a similar set of loops around the bound nucleotide, namely the P-loop, Switch-I, and Switch-II. The magnesium ion bound to the nucleotide displays a similar coordination shell, being in contact with a residues (serine or threonine) from the P-loop and Switch-I. In Fig. 7 we show the structure of myosin (26), kinesin (27), ras (28), and rhoA (29) in what is believed to be a near transition state conformation for nucleotide hydrolysis – the protein is bound to ADP or GDP, with the phosphate replaced by 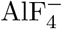 or AlF_3_. After aligning the structures of the four P-loops, the location of Switch-I, Switch-II, and the ligands nearly superposed. In particular, the residues in direct contact with Mg^2+^ show the same position in the binding pocket. With these observations, we surmise that, given the similarity of the binding pocket structure, other ATPases or GTPases may also require the four-fold hydration of magnesium in order to release the phosphate. Thus, the occurrence of step-wise hydration of Mg^2+^ is a pre-requisite for phosphate release in a number of machines, which are unrelated by function and share little sequence or structural similarity.

**Figure 7:**
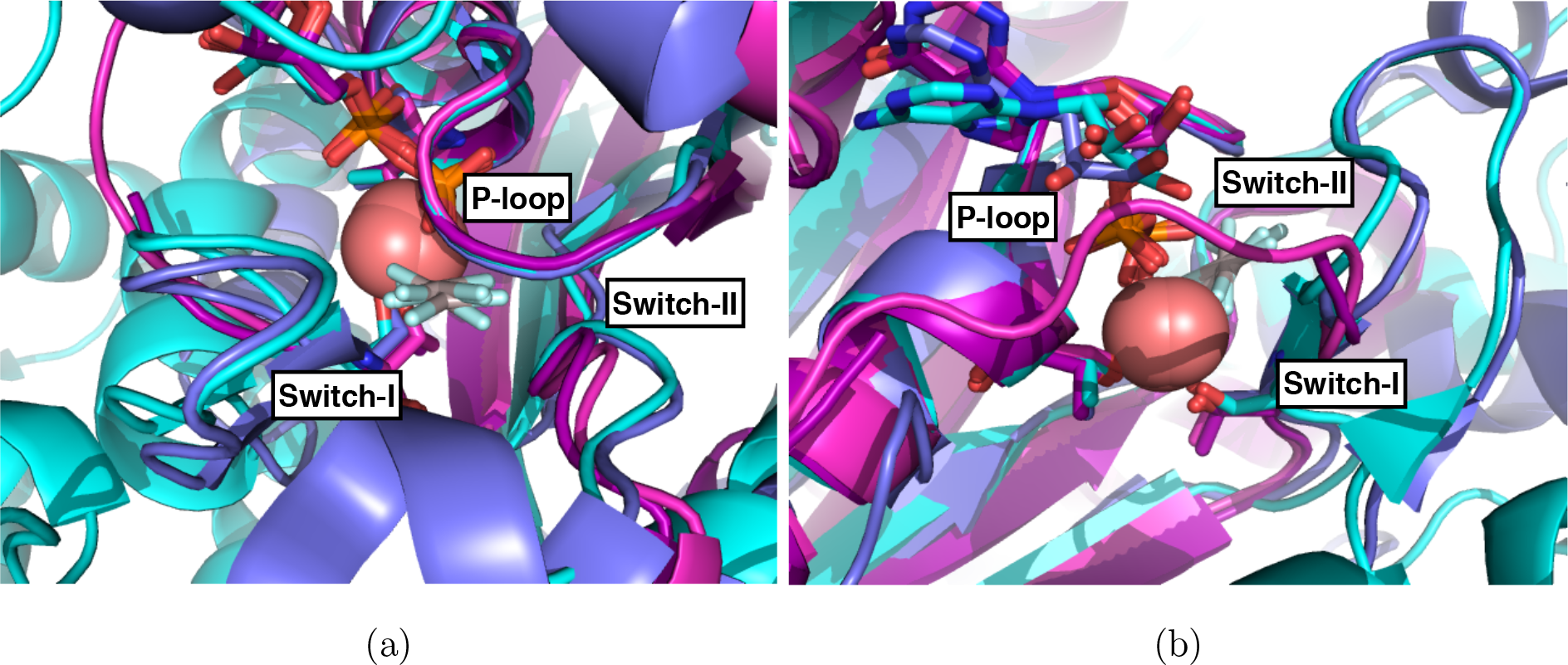
Comparison between the nucleotide binding pockets of molecular motors and G-proteins. In both panels the cartoon represents myosin (cyan), kinesin (blue), ras-rasGAP (magenta), and rho-rhoGAP (purple). In sticks we show the nucleotide (ADP for motors, GDP for G-proteins) and the ligand 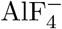 (AlF_3_ for ras) which is believed to mimic the transition state (or an intermediate) for hydrolysis. As sticks, we also show two amino acids interacting with Mg^2+^: T158 (myosin), T92 (kinesin), T19 (rhoA), and S17 (ras) at the end of the P-loop, and S204 (myosin), S202 (kinesin), T37 (rhoA), and T35 (ras) in Switch-I. The red spheres indicate the location of Mg^2+^. The P-loops of the four proteins were aligned with each other using PyMol (43). The nucleotides, cation, ligand, and switches nearly superposed. The PDB used were 4ANJ for myosin (26), 4HNA for kinesin (27), whereas for the G-proteins we used 1WQ1 and 1TX4 for ras (28) and rhoA (29), respectively.

## Conclusions

Extensive atomic detailed molecular dynamics simulations were used to establish that release of phosphate from the nucleotide binding pocket of a myosin motor occurs only after step-wise hydration of Mg^2+^ by four water molecules. This crucial event occurs in many biological machines, and is associated with crucial structural transitions. This prompted us to assess if the mechanism of phosphate release is general. Based on structural comparisons of nucleotide binding pocket in other machines, that bear little relation to the structures or functions of myosin motors, we propose that mechanism for phosphate release is shared among a large class of ATPases and GTPases. This class covers a large number of biological machines. In all the trajectories from release of phosphate was observed, we discovered that the hydration of the first coordination shell of magnesium by four water molecules precedes the expulsion of P_i_. Our finding, which could be universally employed by nature, awaits experimental tests.

## Acknowledgments

We thank Prof. Yale E. Goldman for referring us to the oxygen exchange studies on myosin. We also thank Dr. D. Chakraborty, Dr. A. Kumar, and Dr. S. Shin for a critical reading of the manuscript. This work was supported in part by the National Science Foundation (CHE 19-00033) and the Collie-Welch Chair administered through the Welch Foundation (F-0019). Anton 2 computer time was provided by the Pittsburgh Supercomputing Center (PSC) through Grant R01GM116961 from the National Institutes of Health. The Anton 2 machine at PSC was generously made available by D.E. Shaw Research.

## Methods

### Crystal structure

We started from the crystal structure in PDB 4PJM (6), which was obtained after soaking the crystal in a phosphate bath. The structure indicates that the P_i_ and a few residues are found in two alternative conformations labeled A and B. The simulations employed conformation B, in which the P_i_ and R136 (at the interface between N-terminal region and the converter) are closer to the spatial arrangement of P_i_ in structure 4PFP. In addition, the hydrogen bond between S153 and P_i_, which is formed in structure 4PFP, is broken if the B conformation of S153 is chosen. This likely weakens the phosphate interaction in the nucleotide binding site and may facilitate its release. The water molecules identified in the crystal structure were maintained as a part of the model. In the crystal structure 4PJM, four segments of the myosin motor were not resolved (175-179, 397-406, 565-566, 623-637). We modeled them using Swiss-PDBViewer (30). Because they are far from the nucleotide binding site, it is unlikely that their precise structure significantly affects the release of P_i_. To establish the protonation state of the histidines, we used ProPKa (31, 32).

### Force Fields

The simulations were performed using the CHARMM36 (33) force field for the protein, ADP, K^+^, Cl^−^, and Mg^2+^, with the exception of the simulations in which we used the 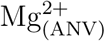 cation from (17), and four simulations performed with reduced magnesium of charge of 1.5. Water molecules were modeled using the TIP3P force field (34). For the phosphate we used 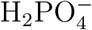, which is the protonation state for the phosphate at neutral pH in low ionic-strength solutions (35), and it has been used as a product in a quantum-mechanical simulation of myosin-catalyzed ATP hydrolysis (18, 19). Moreover, it was proposed that 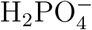 is the form of P_i_ during release (36), which is in agreement with simulations showing that a singly-protonated phosphate is expelled more rapidly than 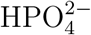 – a result obtained in the simulations of actin (37) and F1 ATPase (13). To generate the force field parameters for 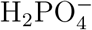 we used CGenFF (38, 39). The motor protein was solvated in a box of water using VMD (40), leaving ⪆1.5nm between the protein and each side of the box. The system was neutralized by 2 atoms of K^+^, and ≈ 100 K^+^ and Cl^−^(corresponding to ≈0.15M of KCl) were also added to approximately mimic the cytosolic salinity. The size of the box, number of water molecules and of ions vary slightly in the different simulations.

### Energy Minimization and equilibration

Before and after solvation, the energy of the system was minimized using NAMD (41) by fixing the position of all the atoms solved by X-ray with the exception of the oxygens of P_i_. After the second round of minimization, we performed a short pre-equilibration of 100ps in which the temperature was slowly raised from 0K to ≈300K. This equilibration, as well as the rest of the simulations, were performed by restraining the position of the *α*-carbons of the residues V643-N652 to avoid rigid body motion of the protein. The tethering potential is harmonic, with spring constant 1kcal/(mol Å^2^). Residues V643-N652 constitute part of a helix in the core of the motor which does not change if the structures of PrePS, P_i_R, and nucleotide-free states are superposed. For this reason these residues were chosen as they are likely not involved in large structural changes during the power stroke (see Fig. S1). After the initial, short equilibration, we ran another equilibration for 7.5ns at a constant pressure (1Atm) and temperature (300K). In order to get consistent results, the last part of the equilibration was always performed using NAMD 2.11 or 2.12, although previous steps of the equilibration were performed using earlier versions of NAMD (2.10) for setup RA-1 (see Table S1). Overall, we found that there is good agreement between the energy and volume sampled during the pressure equilibration performed with NAMD and during the Anton 2 simulations. We excluded from the analysis a few previous simulations performed with Anton (3) and Anton 2 (2) in which the energy, pressure, and volume of the box obtained during equilibration did not match the values reported during sampling. This was a cautious choice because the observables monitored in the excluded simulations did not show any appreciable difference from those discussed here. Unless specified otherwise, the equilibrated structures are referred to as the initial structures.

### Simulations on Anton 2 and data analyses

We carried out simulations on Anton 2. The list of simulations and other technical details are described in the SI. The equations of motion were integrated with the Multigrator algorithm (42) the temperature used is 300K, and the pressure 1Atm. The standard settings were employed to perform the simulations. The energy, pressure, and volume were kept constant (within fluctuations) throughout the system. The extended energy (*E*_*X*_) associated with the Multigrator algorithm underwent a drift. We compared Δ*E*_*X*_ = |*E*_*X*_(*t*) − *E*_*X*_(0)|/|*E*_*X*_(0)| with the results from (42). We found that Δ*E*_*X*_ was about twice the value from (42) if a time step of 2.5fs was used, and about 10 times smaller if the timestep was 2fs. Hence, in most simulations, we used 2fs as a time step. Multiple trajectories (See Table 1 in the SI) using the same initial structures were generated using a different set of initial velocities. For data analyses, we considered only conformations that are 24ns away from each other. A finer temporal resolution is taken into account exclusively to monitor the trajectory of phosphate release. The data analyses was carried out with VMD (40) and PyMol (43).

