## Supplementary Text for "Step-wise Hydration of Magnesium by Four Water Molecules Precedes Phosphate Release in a Myosin Motor"

M.L. Mugnai[1] and D. Thirumalai[1]

[1]Department of Chemistry, The University of Texas at Austin, Austin, TX 78712

October 23, 2019

**List of simulations.** We conducted 24 simulations starting from the same PDB structure (4PJM), and following similar equilibration procedures described in the following paragraphs. The list of simulations is in Table S1. In 4 simulations, we began with the $P_i$ in conformation CA("Crystallographic Alignment", which is explained in the main text) and using the magnesium force field from CHARMM36 (2). We refer to the initial structure as CA-1. Other 5 simulations were performed from a similar initial structure, but with an optimized force field for $Mg^{2+}$ taken from (1), which was calibrated to yield a reasonable value for water exchange times. We refer to this magnesium as $Mg^{2+}_{(ANV)}$, and to the initial structure in these simulations as CA-2. We ran 10 simulations starting from four different initial structures of the RA ("Rotated Alignment", see main text) type, and using the parameters for $Mg^{2+}$ given in CHARMM36 . In order to expedite hydration of $Mg^{2+}$ in 4 simulations we created an artificial magnesium of charge 1.5; to ensure charge neutrality of the box the 100 chlorine ions were assigned a charge of $-0.995$. Also, we performed one simulation restraining the position of the $C_\alpha$ of all the residues to their initial position (last row in Table S1). The only two alpha carbons allowed to move freely were those of the residues that are in contact with $Mg^{2+}$ (T158 and S204).

Finally, we also discussed a short simulation in which starting from the RA orientation and using the $Mg^{2+}_{(ANV)}$ potential for magnesium we observe a quick transition to the CA geometry. The transition occurred during the equilibration phase, so it is not reported in Table S1, in which only post-equilibration trajectories are discussed. We performed these simulations not only to ensure that our major conclusion, announced in the title, is robust, but also in order to observe $P_i$ release in the limited time scales accessible to standard simulations.

### The coordination shell of Magnesium

**CA, CHARMM force field for $Mg^{2+}$.** In Fig. S2 we show the coordination of $Mg^{2+}$ in the trajectories obtained with CHARMM36 (2) force field for $Mg^{2+}$ starting from the CA conformation. One of these simulations was shown in Fig. 3 of the main text and it is not reported here. During these trajectories the number of water molecules bound to $Mg^{2+}$ is either 2 or 3, and the phosphate is not released on the timescale of the simulations.

**CA, ANV force field for $Mg^{2+}$.** In Fig. S3 we show the coordination of $Mg^{2+}$ in the trajectories carried out with the force field $Mg^{2+}_{(ANV)}$ for the cation (1), starting from the CA conformation. Occasionally, we observe that the number of water molecules in the first coordination shell of the cation is four, but the phosphate is not released during the simulations.

**RA, CHARMM force field for $Mg^{2+}$.** In Fig. S4 we show the coordination of $Mg^{2+}$ in the trajectories generated using CHARMM36 (2) force field for $Mg^{2+}$ starting from the RA conformation. A segment of one of these simulations is shown in Fig. 4

Table S1: List of the simulations performed. The initial structure for each simulation (that is, after the initial fast equilibration) is in the first column, and the length of the trajectory is in the second column. Columns 3-5 describe information about $Mg^{2+}$: force field used (third column), list of molecules that constitute the solvation shell at the beginning (fourth column) and at the end of the simulation (fifth column). In the last column we report whether $P_i$ has escaped. The initial structures are shown in Fig. 2b (CA-1), Fig. 2c (CA-2), and Fig. 2d (RA-3). With ANV we refer to the force field in (1), and with C36 we indicate CHARMM36 (2). In describing the solvation shell of $Mg^{2+}$ (column 4 and 5), we use W to indicate water, D for ADP, T for T158 side chain, S for S204, and $Cl^-$ is a chlorine ion.

| Initial structure | $\approx t_{end}$ | $Mg^{2+}$ | $Mg^{2+}(t=0)$ | $Mg^{2+}(t_{end})$ | Release |
|---|---|---|---|---|---|
| CA-1 | 8.4$\mu$s | C36 | 2W,D,$P_i$,T,S | 3W,2D,$P_i$ | No |
| CA-1 | 5.4$\mu$s | C36 | 2W,D,$P_i$,T,S | 3W,2D,$P_i$ | No |
| CA-2 | 4.8$\mu$s | ANV | 2W,D,$P_i$,T,S | 3W,2D,$P_i$ | No |
| CA-1 | 4.8$\mu$s | C36 | 2W,D,$P_i$,T,S | 3W,2D,$P_i$ | No |
| CA-1 | 4.8$\mu$s | C36 | 2W,D,$P_i$,T,S | 3W,2D,$P_i$ | No |
| CA-2 | 7.2$\mu$s | ANV | 2W,D,$P_i$,T,S | 2W,3D,$P_i$ | No |
| CA-2 | 4.8$\mu$s | ANV | 2W,D,$P_i$,T,S | 3W,2D,$P_i$ | No |
| CA-2 | 4.8$\mu$s | ANV | 2W,D,$P_i$,T,S | 3W,2D,$P_i$ | No |
| CA-2 | 4.6$\mu$s | ANV | 2W,D,$P_i$,T,S | 3W,2D,$P_i$ | No |
| RA-1 | 4.8$\mu$s | C36 | 2W,3D,T | **4W**,2D | Yes |
| RA-1 | 4.8$\mu$s | C36 | 2W,3D,T | **4W**,2D | Yes |
| RA-2 | 4.8$\mu$s | C36 | 2W,3D,T | **4W**,2D | Yes |
| RA-2 | 4.8$\mu$s | C36 | 2W,3D,T | 3W,3D | No |
| RA-2 | 4.8$\mu$s | C36 | 2W,3D,T | **4W**,2D | Yes |
| RA-2 | 4.8$\mu$s | C36 | 2W,3D,T | 2W,3D,T | No |
| RA-3 | 4.8$\mu$s | C36 | 2W,3D,T | **4W**,2D | Yes |
| RA-3 | 4.8$\mu$s | C36 | 2W,3D,T | 3W,2D,$P_i$ | No |
| RA-3 | 4.8$\mu$s | C36 | 2W,3D,T | **4W**,2D | Yes |
| RA-3 | 4.8$\mu$s | C36 | 2W,3D,T | 3W,2D,T | No |
| CA-3 | 4.8$\mu$s | 1.5 | 2W,2D,$P_i$ | 2W,2D,$P_i$ | No |
| CA-3 | 4.8$\mu$s | 1.5 | 2W,2D,$P_i$ | 2W,2D,$P_i$ | No |
| CA-3 | 13.2$\mu$s | 1.5 | 2W,2D,$P_i$ | 2D,$P_i$,$Cl^-$ | No |
| CA-3 | 4.8$\mu$s | 1.5 | 2W,2D,$P_i$ | 2W,2D,$P_i$ | No |
| CA-1 | 2.4$\mu$s | C36 | 2W,D,$P_i$,T,S | 2W,2D,$P_i$ | No |

of the main text. During those trajectories in which the phosphate is not released, the number of water molecules bound to $Mg^{2+}$ is four only for a short period of time in one trajectory.

### The Rotation of the Converter

During the simulations the converter moves away from the pre-stroke pose, though it never reaches the conformation observed in the post-stroke state. We show in Fig. S7 the root mean squared deviation (RMSD) of the converter computed by aligning the rest of the motor domain (upper figure in each panel) or the converter (lower figure). The analysis indicates that the changes in RMSD observed when the motor protein is aligned are largely due to the rigid body motion of the converter, whose structure is maintained in the vicinity of the crystallographic conformation during the course of the simulation.

### The Movement of the N-Terminal Beta Barrel

We report also an unexpected movement of the N-terminal $\beta$ barrel (NTBB) observed in a few trajectories. In Fig. S8 we show that, as well as in the case of the movement of the converter, the majority of the RMSD from the initial conformation is associated with the rigid-body motion of the NTBB, whose structure, instead, appears to remain in the vicinity of the crystallographic conformation.

### The Movement of ADP

In the majority of the simulations, the ADP maintains the contacts with the protein. On the other hand, it appears that the presence of $P_i$ in the RA conformation favors some movement of ADP in the nucleotide binding site. In Fig. S9, we report a contact between ADP and the residue N98, and show how in those simulations in which the phosphate is rotated and not released the interaction with the myosin is weakened.

### Movies

The movie PiRelease_AllMotor.mpg (Supplementary Movie 1) shows the whole myosin VI during a trajectory in which the phosphate is released. From the visualization angle, the movement of the converter and the NTBB are clear.

The movie PiRelease_Pocket.mpg (Supplementary Movie 2) focuses on the nucleotide binding pocket. ADP, magnesium, and phosphate are shown as spheres, water, T158 and S204 as sticks. After the fourth water molecule binds the cation the phosphate is

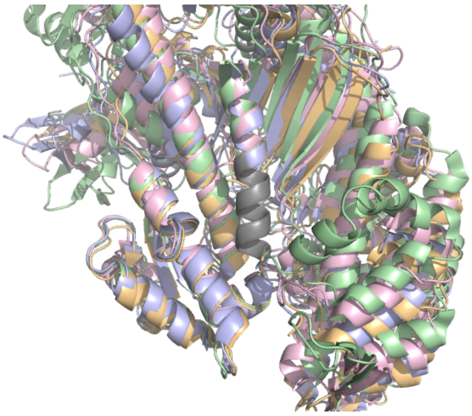

Figure S1: Residues restrained to reduce the rigid body motion of myosin VI. The restrained residues V643-N652, shown in grey, are used to align the ADP-bound [PDB 6BNW (3), shown in pink], PiR [4PJM (4), shown in orange], rigor [2BKI (5), shown in green], and Pre-PS [4ANJ (6), shown in blue] myosin VI, all shown in the figure. Because the grey helix is well aligned in all of these conformations, we chose to restrain the $\alpha$-carbons of this helix in order to remove the rigid body motion of the protein. The rationale is that if the conformation of the helix has not changed in these different structures, by keeping the helix fixed we are not preventing some relevant conformational transition.

expelled. In this trajectory only, we observe that the anion returns in the binding pocket, is released again, and then finally re-enters.

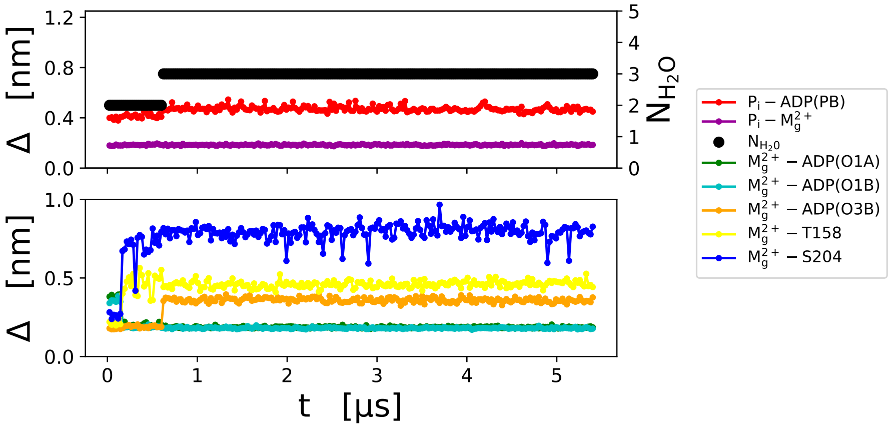


(a)

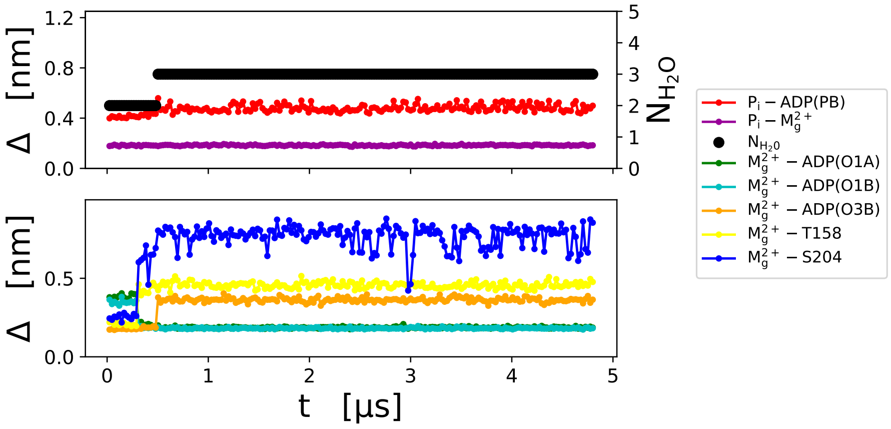


(b)

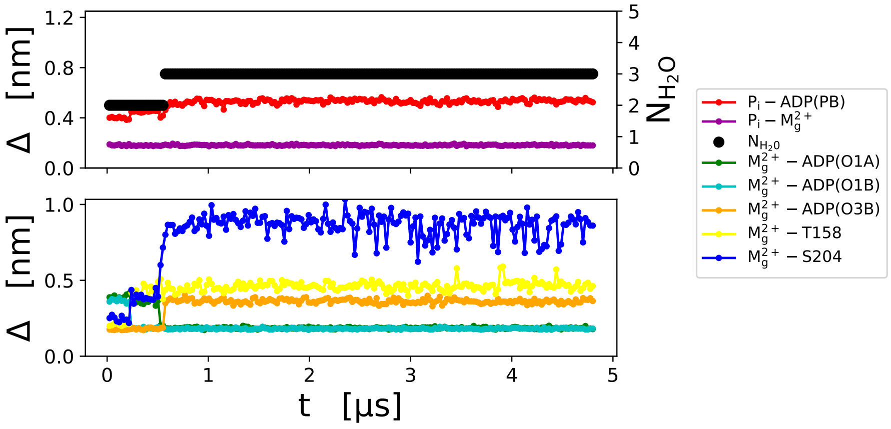


(c)

Figure S2: Coordination of magnesium during a simulation carried out from CA initial conformation with CHARMM $Mg^{2+}$. Time-dependent distances of atoms engaged in the first solvation shell of $Mg^{2+}$ are shown. The color code is reported in the legend and discussed in the caption of Fig. 3 in the main text. The three panels show the results from different simulations.

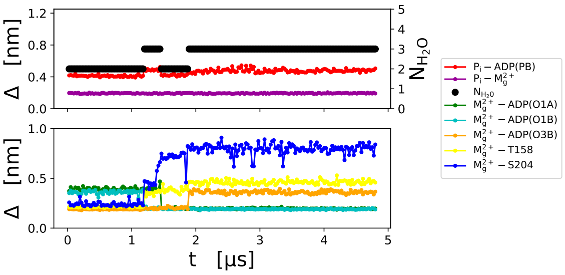


(a)

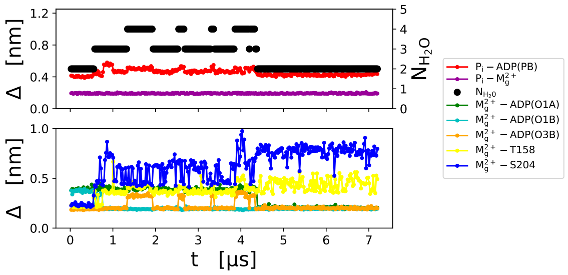


(b)

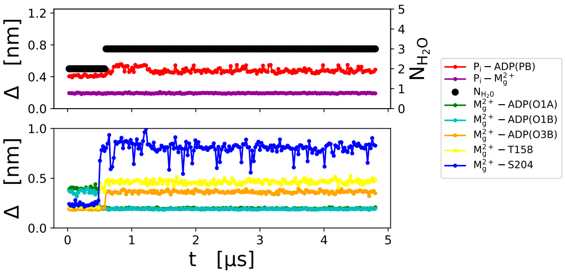


(c)

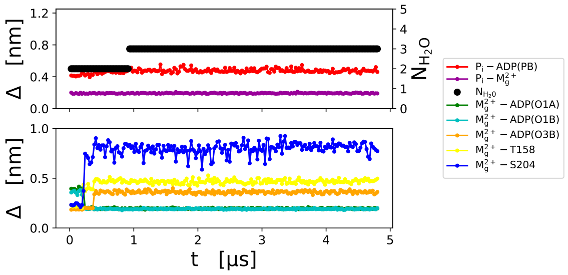


(d)

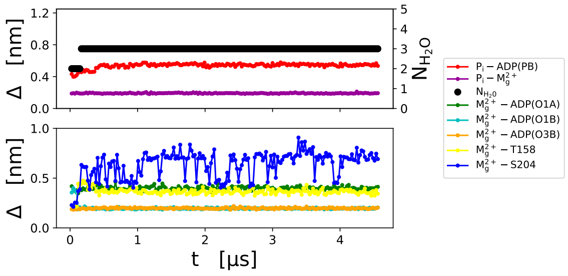


(e)

Figure S3: Coordination of magnesium during a simulation carried out from the CA initial conformation with $Mg^{2+}_{(ANV)}$ (1), with parameters developed to account for reasonable water exchange rate of isolated $Mg^{2+}$ in pure water. Time-dependent distances of atoms engaged in the first solvation shell of $Mg^{2+}$. The color code is the same as in Fig. 3. The five panels show results from different trajectories.

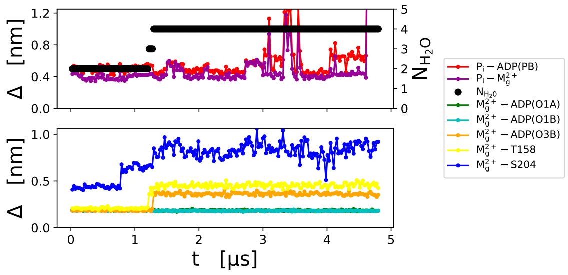


(a)

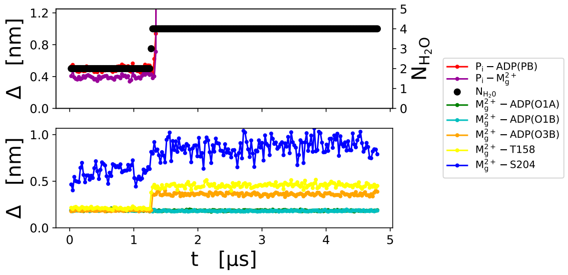

(b)

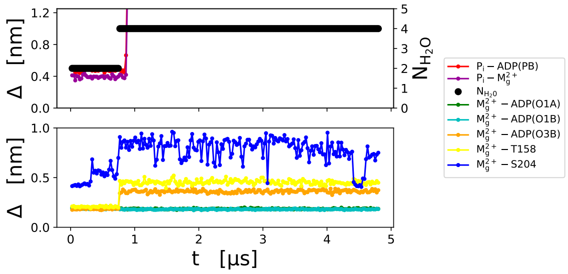

(c)

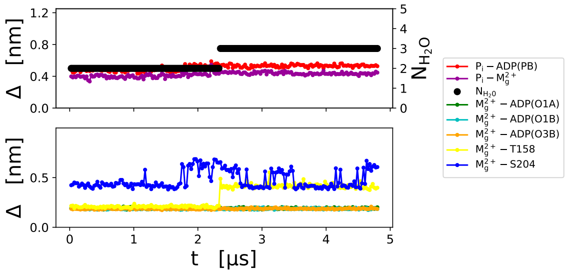

(d)

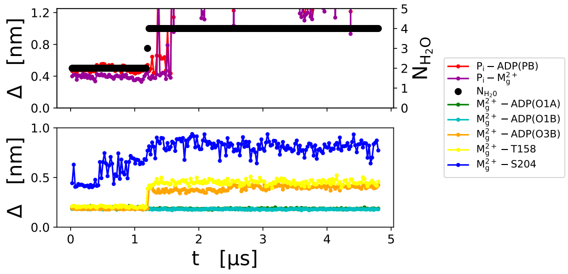

(e)

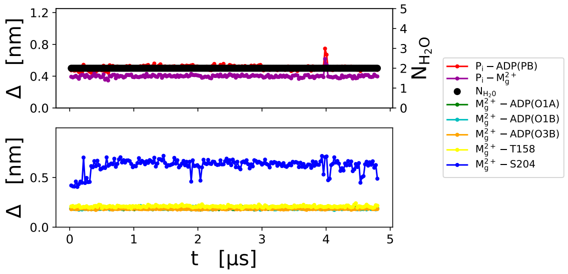

(f)

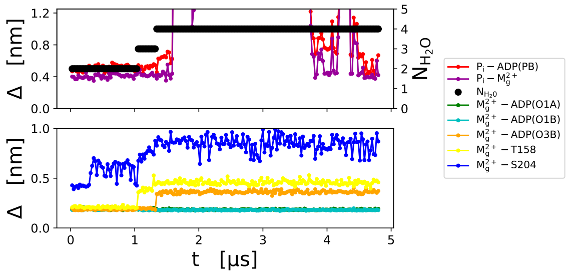

(g)

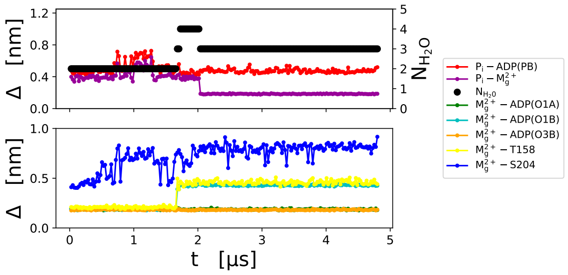

(h)

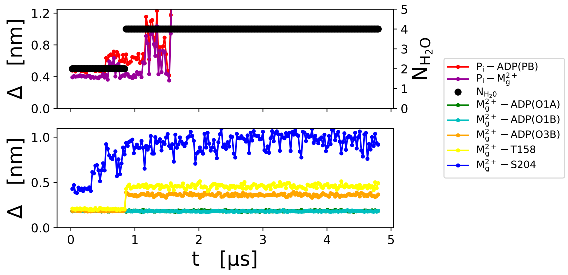

(i)

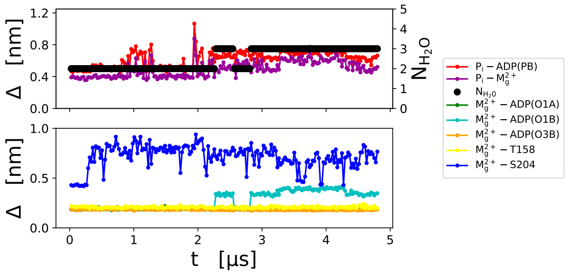

(j)

Figure S4: Coordination of magnesium during a simulation carried out from the RA initial conformation. Time-dependent distances of atoms engaged in the first solvation shell of $Mg^{2+}$. The color code is the same of Fig. 3. The ten panels show results from different trajectories.

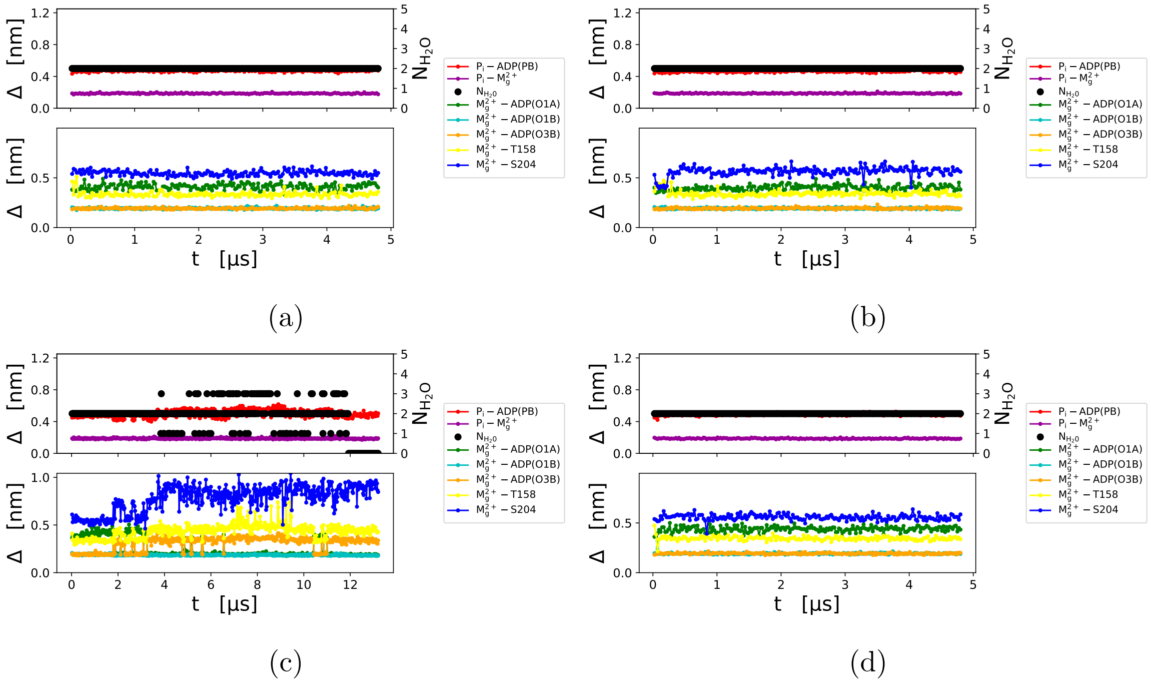


Figure S5: Coordination of magnesium during a simulation carried out with the weaker magnesium cation. Time-dependent distances of atoms engaged in the first solvation shell of $Mg^{2+}$. The color code is the same of Fig. 3. The four panels show results from different trajectories.

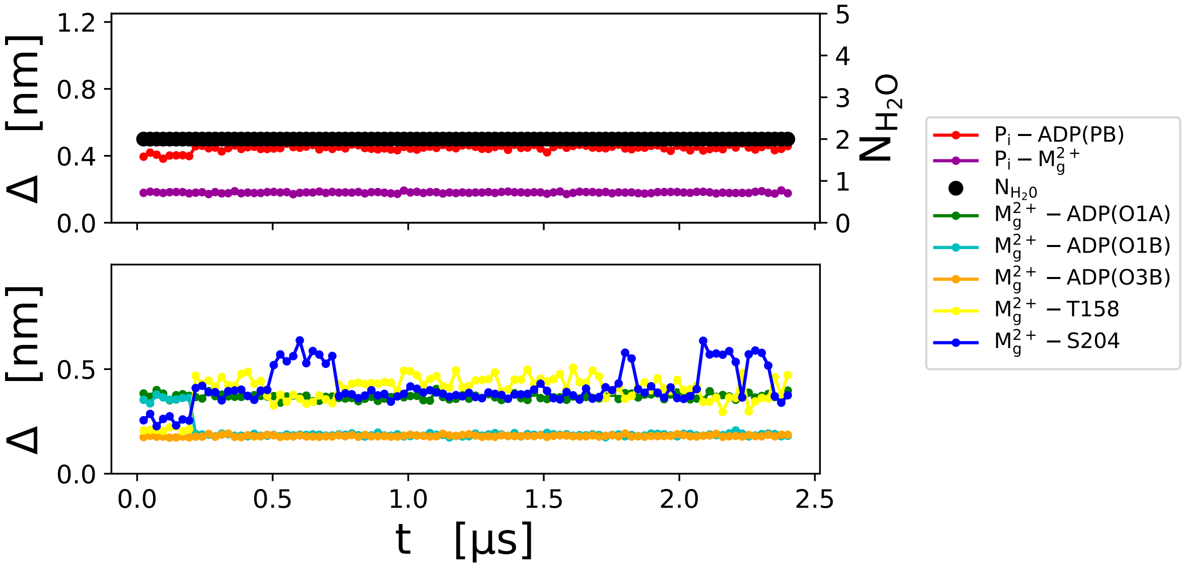


Figure S6: Coordination of magnesium during a simulation carried out restraining the alpha carbons of all residues, with the exception of T158 and S204.

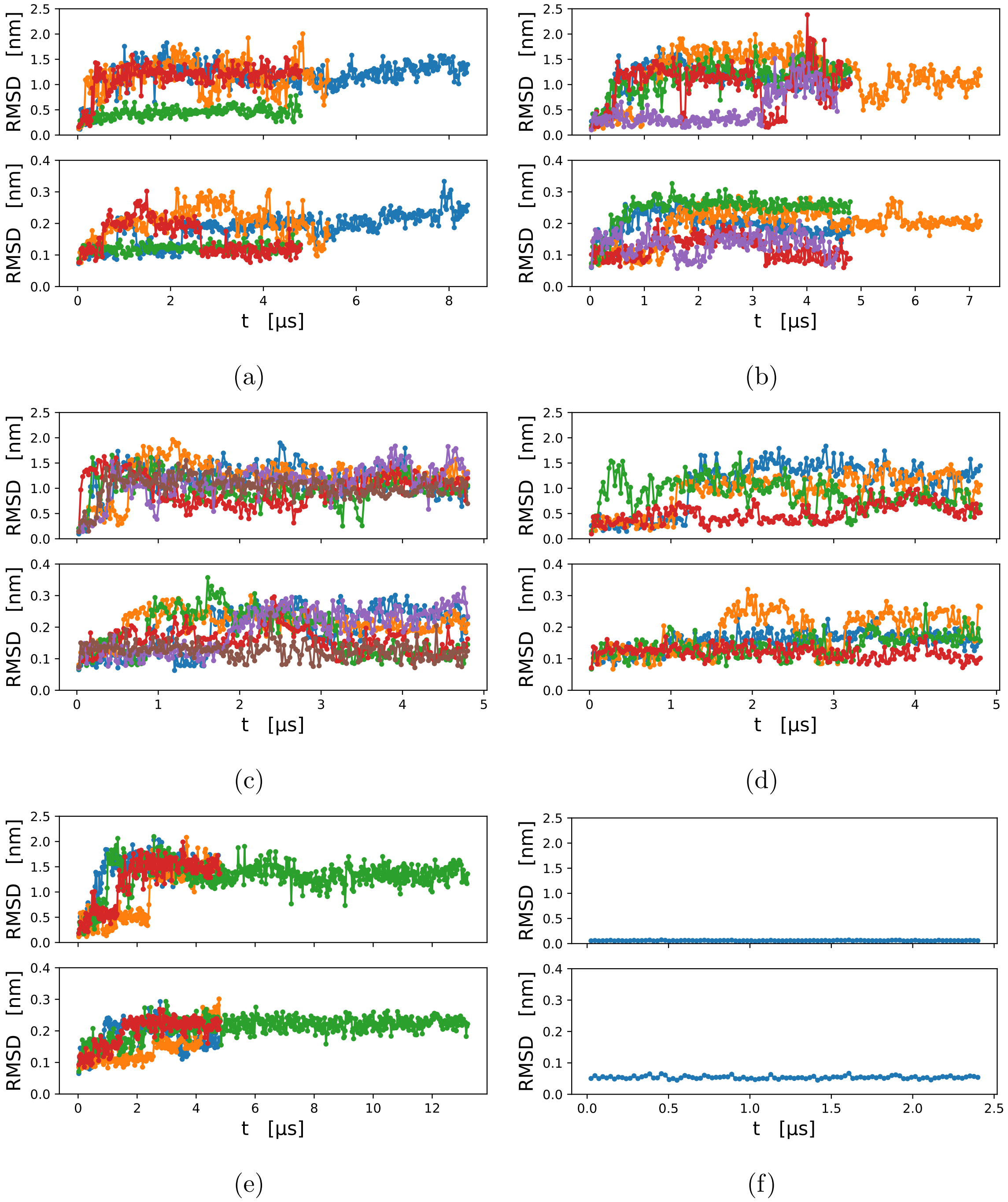


Figure S7: Movement of the converter. There are 5 panels each made of two figures sharing the x-axis, where simulation time is shown. For each panel, the upper figure shows the RMSD of the converter with respect to the initial conformation as a function of time when myosin VI (residues 51-705) was aligned with the initial structure and the RMSD of the converter (706-773) was computed. The lower figure shows the case in which the converter was aligned to the initial structure. Note the difference in the scale between the upper and lower figures. (a) Simulations carried out with the CHARMM $Mg^{2+}$ starting with the CA conformation. (b) Results obtained with $Mg^{2+}_{(ANV)}$. (c) Trajectories starting from the RA conformation esulting in the release of phosphate. (d) Simulations initialized with the RA conformation, during which the phosphate remains in the nucleotide binding site. (e) Simulations carried out with the weakly-charged magnesium. (f) Simulations in which the alpha carbons of all residues (with the exception of T158 and S204) were restrained.

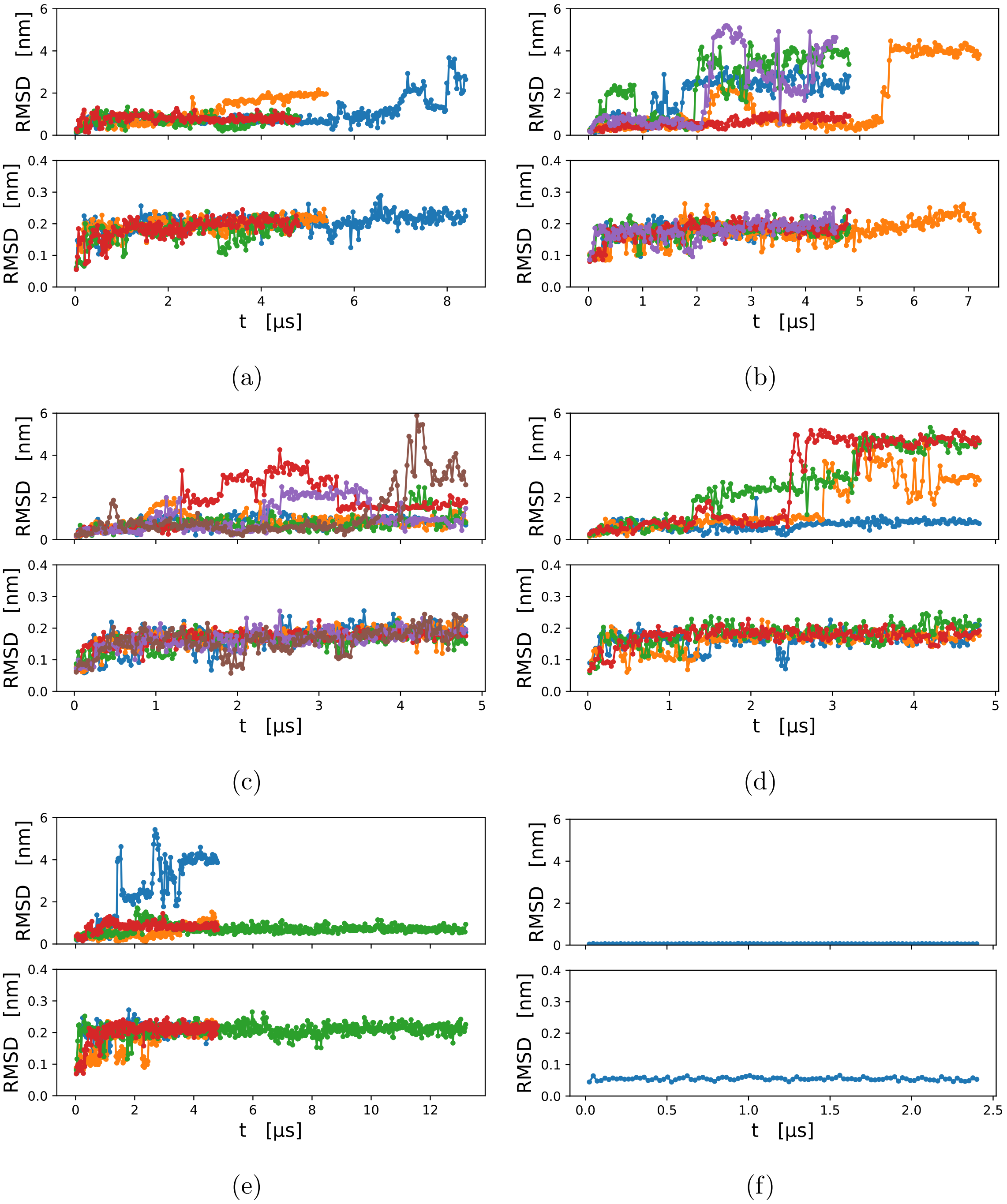


Figure S8: Movement of the NTBB. For each panel, the upper figure shows the RMSD of the NTBB with respect to the initial conformation as a function of time when the protein (residues 51-705) was aligned to the initial structure and the RMSD of the NTBB (5-50) was computed. The lower figure shows the case in which the NTBB was aligned to the initial structure. The panels are the same as in Fig. S7.

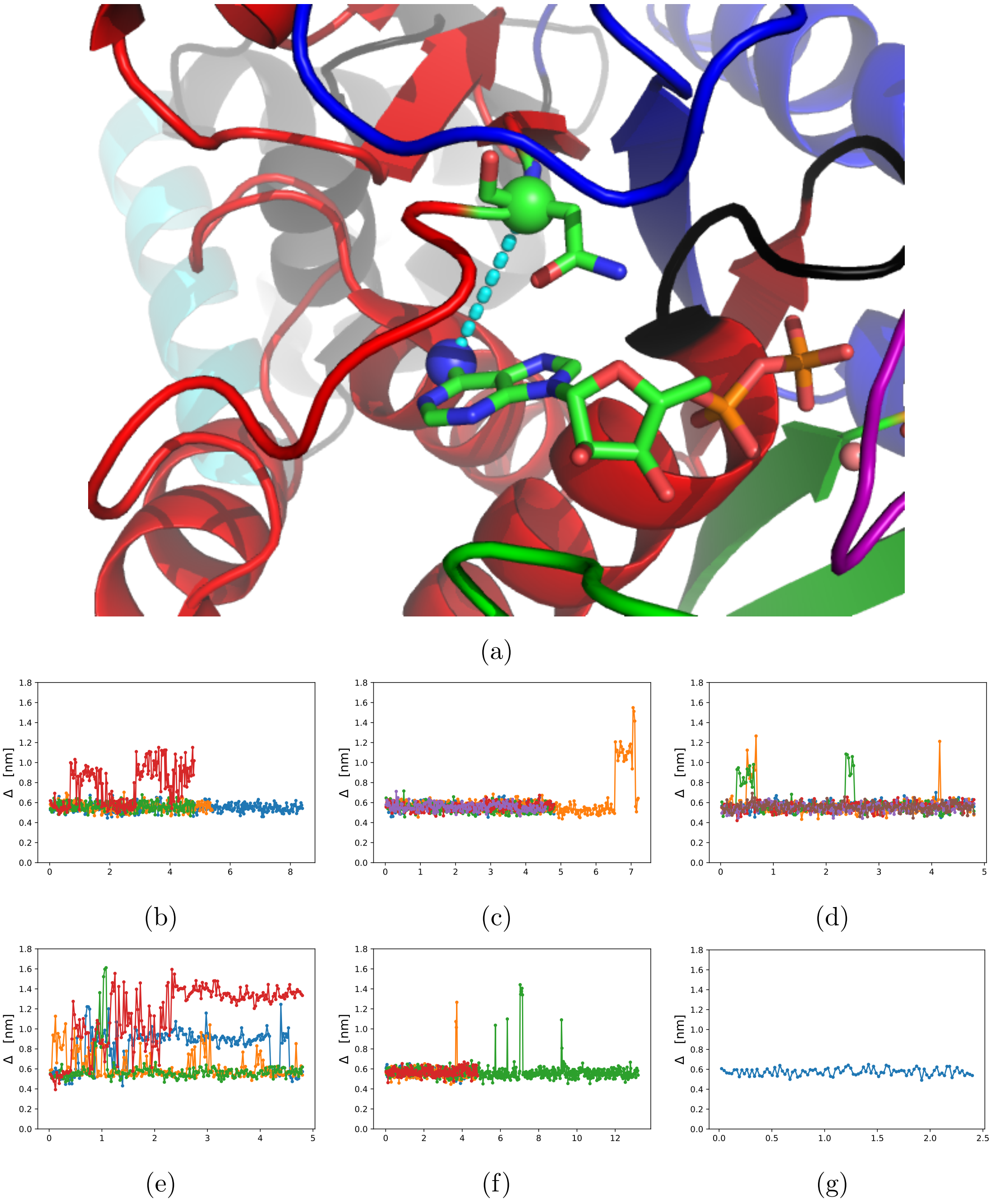


Figure S9: Movement of the ADP. We show the distance between the nitrogen N6 of ADP and the $\alpha$ carbon of residue N98. Panel (a): the cyan, dashed line shows the monitored distance between the nitrogen N6 of ADP (blue sphere) and the alpha carbon of N98 (green sphere). Panels (b-c): time-dependent results for simulations initialized in CA with CHARMM36 force field for magnesium (b) or ANV parameters (c). Panels (d-e): results for simulations with RA initial alignment ending in the expulsion of the phosphate (d) or without $P_i$ escape (e). Panels (f-g): time-traces when a weakly-charged magnesium was used (f), or with harmonic restraints applied to all the alpha carbons (with the exception of T158 and S204) (g).
